# The Rise and Fall of Slow Wave Tides: Vacillations of Slow Wave/Spindle Coupling Shift the Composition of Slow Wave Activity Through Sleep Cycles in Accordance with Depth of Sleep

**DOI:** 10.1101/2022.03.22.485383

**Authors:** Brice V. McConnell, Eugene Kronberg, Lindsey M. Medenblik, Vitaly O. Kheyfets, Alberto R. Ramos, Stefan H. Sillau, Rachelle L. Pulver, Brianne M. Bettcher

## Abstract

Slow wave activity (SWA) during sleep is associated with synaptic regulation and memory processing functions. Each cycle of non-rapid-eye-movement (NREM) sleep demonstrates a waxing and waning amount of SWA during the transitions between stages N2 and N3 sleep, and the deeper N3 sleep is associated with an increased density of SWA. Further, SWA is an amalgam of different types of slow waves, each identifiable by their temporal coupling to spindle subtypes with distinct physiological features. The objectives of this study were to better understand the neurobiological properties that distinguish different slow wave and spindle subtypes, and to examine the composition of SWA across cycles of NREM sleep. We further sought to explore changes in the composition of NREM cycles that occur among aging adults. To address these goals, we analyzed subsets of data from two well-characterized cohorts of healthy adults: 1) The DREAMS Subjects Database (n=20), and 2) The Cleveland Family Study (n=60). Our analyses indicate that slow wave/spindle coupled events can be characterized as frontal versus central in their relative distribution between electroencephalography (EEG) channels. The frontal predominant slow waves are identifiable by their coupling to late-fast spindles and occur more frequently during stage N3 sleep. Conversely, the central-associated slow waves are identified by coupling to early-fast spindles and favor occurrence during stage N2 sleep. Together, both types of slow wave/spindle coupled events form the composite of SWA, and their relative contribution to the SWA rises and falls across cycles of NREM sleep in accordance with depth of sleep. Exploratory analyses indicated that older adults produce a different composition of SWA, with a shift toward the N3, frontal subtype, which becomes increasingly predominant during cycles of NREM sleep. Overall, these data demonstrate that subtypes of slow wave/spindle events have distinct cortical propagation patterns and differ in their distribution across lighter versus deeper NREM sleep. Future efforts to understand how slow wave sleep and slow wave/spindle coupling impact memory performance and neurological disease may benefit from examining the composition of SWA to avoid potential confounds that may occur when comparing dissimilar neurophysiological events.

## Intro

The slow waves of non-rapid-eye-movement (NREM) sleep are hypothesized to coordinate synaptic remodeling and sleep-dependent memory processes^1,2^. Studies indicate that loss of normal slow wave physiology is related to cognitive impairment and pathophysiological changes of Alzheimer’s disease, suggesting that slow waves may have particular relevance to pathological aging processes^3–8^. Efforts to understand the neurobiology of slow waves have noted that slow wave activity (SWA) is not uniform in nature and is instead a heterogeneous composition of different subtypes of slow waves. Within surface EEG, SWA has been separated into subcomponents by several slow wave characteristics, including frequency’, local versus global occurrence, and frontal versus central localization.

Slow waves can also be conceptualized as discrete neuronal communication events that mark time windows of hippocampal activity and reactivation of specific neuronal patterns. During these slow wave time windows, neuronal patterns recapitulate the original sequences of memory tracings that first formed during wake within the hippocampi and association cortexes^14–16^. In this mechanistic framework, the EEG signals associated with slow waves, termed coupled sleep spindles, are additional components of a “memory replay mode” that occurs during NREM sleep^16,17^. Intracranial recordings further support a distinction between subtypes of slow waves based on their temporal coupling with anterior versus posterior hippocampal activity, and the relative contribution of these distinct slow wave and coupled spindle components to SWA differs between stages N2 and N3 sleep^18,19^. Within surface EEG, two distinct subtypes of slow waves can also be identified by their coupling to unique spindle subtypes, and congruent with intracranial observations, the contribution of each coupled slow wave/spindle subtype to SWA varies between stages N2 and N3 sleep.^20^

Experimental manipulation of SWA-associated memory functions in animal models further supports a physiological distinction between different subtypes of coupled slow wave/spindle events, with one subtype of coupled events promoting memory retention, while another is observed to facilitate weakening of memory representations^21^. Notably, this observed combination of synaptic strengthening and weakening of memory content has been proposed as a possible mechanistic framework for sleep-dependent memory consolidation^22^. Thus, the mixing of different subtypes of coupled slow wave/spindle events during cycles of NREM sleep may impact key and distinct memory consolidation processes underlying selective synaptic remodeling and synaptic homeostasis. Further, given that changes in the depth of sleep between N2 and N3 sleep are associated with shifting compositions of SWA subtypes, these transitions between N2 and N3 sleep across each NREM cycle may create ebbs and flows of different synaptic remodeling activities.

Analyzing the composition of NREM sleep cycles provides a tool to advance our fundamental understanding of sleep-dependent memory consolidation and the proposed neuroprotective aspects of slow wave sleep. Studies of sleep and memory have reported incongruencies regarding slow wave sleep’s memory promoting effects, with some data indicating that slow wave sleep strengthens overnight memory consolidation, while others fail to reproduce these observations^4,23^. Notably, these incongruencies remain poorly explained in a more limited, homogenous, conceptual framework of SWA that does not account for the differences in the composition of slow wave sleep.

Properties of SWA are also proposed to be a possible biomarker of neurodegenerative disease, given that the proposed neuroanatomical components of SWA-associated memory consolidation activities are directly impacted by Alzheimer’s disease processes including beta amyloid and tau accumulation^7,8,24,25^. Disruptions in the specific timing relationships between slow waves and spindles correlate with worsening cognitive performance among aging adults^26,27^, and the presence of Alzheimer’s disease pathology may underly some these timing abnormalities^24^. Thus far, these studies have been limited to analyzing SWA as a mixture of different slow wave and spindle subtypes with unknown composition, and given that the timing aspects of slow wave and spindle coupling are observed to differ between early-fast and late-fast spindles^20^, a more advanced approach to capturing the composition of SWA may provide tools and insights to further delineate the connections between sleep and Alzheimer’s disease processes.

Our study was designed to appraise whether structure and composition of the SWA may be fundamentally different depending on the depth of NREM sleep and time within a sleep session, with an overarching goal to examine how assessment of SWA may vary depending on experimental conditions. We leveraged two existing and well-characterized sleep recording datasets to examine the structure and composition of SWA across the night and within cycles of NREM sleep. We further examined how the composition of SWA through cycles of NREM sleep changes with age, building on prior observations that aging is associated with overall greater late-fast composition of NREM sleep^20^. Here we specifically expanded on this work to examine the temporal composition of SWA among at discrete time intervals, and we measured the relative composition of SWA slow wave/spindle subtypes across the night for each younger and older research participant.

In light of studies suggesting that SWA and the coupling of slow waves to spindles may be an early biomarker of cognitive aging and neurodegenerative disease^7,8,24,28^, efforts to advance these insights and to translate the science to clinical use will need to account for the fundamental changes in SWA composition across cycles of NREM. Here we characterize and illustrate the distinctions in the composition of SWA, and we demonstrate the remarkable variance in the composition of SWA between young adults and aging adults.

## Methods

All data were collected with written informed consent under protocols approved by a local institutional review board at each institution. Participant data was accessed from the DREAMS database^29^ and from the Cleveland Family Study (CFS) via the National Sleep Research Repository.^30-33^ The DREAMS database was originally created to study automated sleep staging techniques among overnight sleep recordings among individuals with no identifiable sleep pathologies including sleep apnea. Recordings from DREAMS are characterized as containing very few EEG artifacts and were included in this database specifically for their recording clarity. We selected the DREAMS database for our sub-analysis based on the quality of the recordings and the availability of frontal and central recording electrodes (these recordings sites are not available within the CFS dataset). Overnight polysomnography was utilized from all of the available 20 normal control participants available within the DREAMS database.

The CFS was originally conducted to study sleep apnea among a well-characterized cohort of 2,284 individuals and we selected this study for our secondary analysis due to the availability of high-quality overnight polysomnography from participants in young adult and older adult age ranges (distinct age ranges are not available within the DREAMS database). Data quality was originally rated by sleep staging experts per standardized study protocol, and we selected recordings for analysis to contain at least 95% visually-scorable EEG data. Individuals from CFS were selected from ages 18-35 for young adults (n=30; median age 24.19 (21.78-25.48 IQR)) and 60-70 for older adults (n=30; median age 65 (62.09-68.13 IQR)), given prior data indicating that these age ranges differ in slow wave/spindle coupling.^20^ To minimize possible effects of sleep apnea, we selected 60 participants within the CFS by lowest oxygen desaturation apneas and hypopneas >= 3% oxygen desaturation or arousal per hour of sleep (AHI) scores within each age desired range (median AHI for young adults = 0.21 (0.15-0.50 IQR)); median AHI for older adults = 4.51 (2.80-9.15 IQR); Table 1).

**Table 1.**
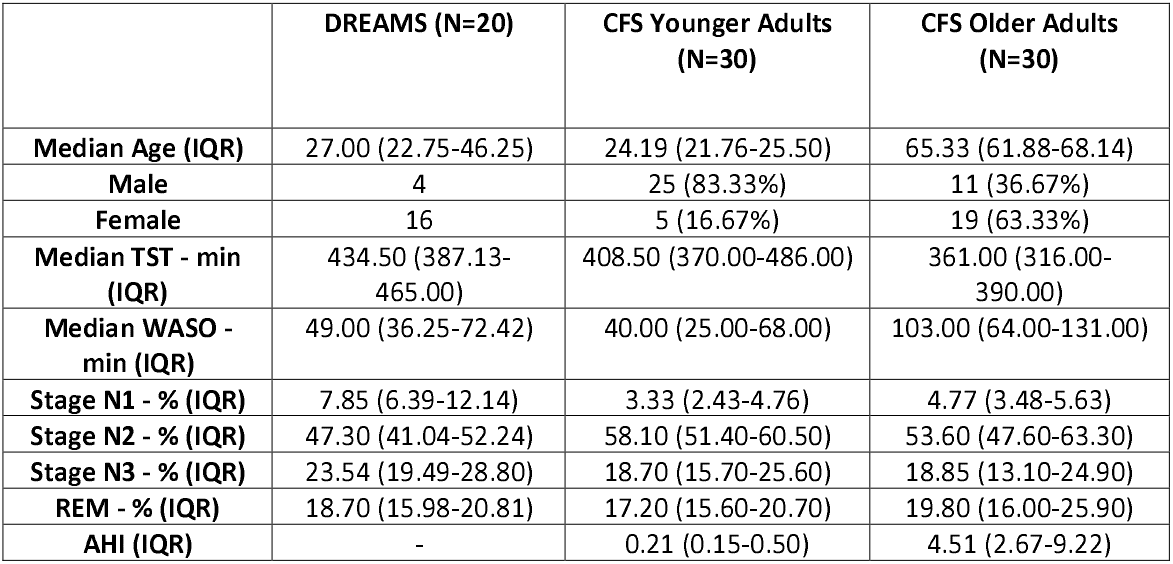
Demographics and sleep characteristics of the study subjects. IQR= Interquartile Range

|  | **DREAMS (N=20)** | **CFS Younger Adults (N=30)** | **CFS Older Adults (N=30)** |
| --- | --- | --- | --- |
| **Median Age (IQR)** | 27.00 (22.75-46.25) | 24.19 (21.76-25.50) | 65.33 (61.88-68.14) |
| **Male** | 4 | 25 (83.33%) | 11 (36.67%) |
| **Female** | 16 | 5 (16.67%) | 19 (63.33%) |
| **Median TST - min (IQR)** | 434.50 (387.13-465.00) | 408.50 (370.00-486.00) | 361.00 (316.00-390.00) |
| **Median WASO - min (IQR)** | 49.00 (36.25-72.42) | 40.00 (25.00-68.00) | 103.00 (64.00-131.00) |
| **Stage N1 - % (IQR)** | 7.85 (6.39-12.14) | 3.33 (2.43-4.76) | 4.77 (3.48-5.63) |
| **Stage N2 - % (IQR)** | 47.30 (41.04-52.24) | 58.10 (51.40-60.50) | 53.60 (47.60-63.30) |
| **Stage N3 - % (IQR)** | 23.54 (19.49-28.80) | 18.70 (15.70-25.60) | 18.85 (13.10-24.90) |
| **REM - % (IQR)** | 18.70 (15.98-20.81) | 17.20 (15.60-20.70) | 19.80 (16.00-25.90) |
| **AHI (IQR)** | - | 0.21 (0.15-0.50) | 4.51 (2.67-9.22) |

### EEG Acquisition

For Dream EEG files, recordings were acquired at the André Vésale hospital (Montigny-le-Tilleu I, Belgium) using a digital 32-channel polygraph (BrainnetTM System of MEDATEC, Brussels, Belgium). At least two EOG channels (P8-A1, P18-A1), three EEG channels (CZ-A1 or C3-A1, FP1-A1 and O1-A1) and one submental EMG channel were recorded at a sampling frequency of 200Hz. Staging was performed by a single expert at the sleep laboratory following AASM criteria.^34^ Similarly for CFS EEG files, laboratory-based polysomnography was performed using a 14-channel Compumedics E-Series System (Abbotsford, Australia). Recordings were created using gold cup electrodes at a sampling rate of 128 Hz (filtered low pass 0.16 Hz and high pass 105 Hz). Electrodes at C3 and C4 were referenced to positions A2 and A1, respectively. Sleep staging was performed by trained technicians at a centralized reading center at Brigham and Women’s Hospital following study protocols and AASM criteria.^34^

### Slow Wave Detection

Slow waves were identified via automated zero-crossing detection as described in McConnell et al.^20^ Briefly, signals for slow wave detection were utilized from electrodes FP1, CZ, and C3 from stages N2 and N3 of slow wave sleep by AASM criteria.^34^ Channels FP1 and CZ were selected for their relatively close positions to slow wave sources described in localization experiments, which demonstrated a frontal source near the left insula and a central source near the cingulate gyrus^13^. Electrode C3 was selected as having been well-characterized in our prior analysis of slow wave and spindle coupling among 582 individuals from CFS^20^.

Epochs with un-scorable data were excluded from analysis. Automated management of high amplitude artifacts was accomplished via exclusion of EEG segments exceeding 900 μV after detrending data with sliding window of three seconds across raw data. EEG data was subsequently detrended and band-pass filtered in a forward and backward direction using a 6th-order Butterworth filter between 0.16-4 Hz. Zero crossings were identified to detect negative and positive half-waves, and slow wave events were identified when the half-wave pairs approximated a frequency range of 0.4 to 4 Hz. Minimal and maximal half-wave amplitudes were measured, and slow waves with both positive and negative maximum amplitudes in the top 50% of all waves were selected for subsequent coupling analysis. An upper threshold of +/- 200 μV for zero crossing pairs was utilized to reduce misidentification of non-slow wave events. A further reduction of false identifications was accomplished by rejecting all zero crossing pairs with peak/trough amplitudes exceeding four standard deviations from the mean min/max zero crossing pair values for each subject.

### Time-Frequency Analysis

Time-frequency wavelet plots of coupled spindles were created via established methods.^20,35,36^ Troughs of each slow wave were centered in 5-second intervals of EEG data and matched to 5-second baseline intervals immediately preceding slow wave events (excluding baseline segments containing slow wave events). Morlet-wavelet transformation (8 cycles) was applied to the unfiltered EEG for slow wave and baseline segments between 4 Hz and 20 Hz in steps of 0.25Hz. The mean of baseline regions was used to normalize the amplitude of the mean Morlet-wavelet transformation of all 5-second peri-slow wave regions.

### Slow Wave Separation by Co-Identification of Spindle Events and Timing Alignment

Spindle event identification was consistent with parameters of prior reports^20,36,37^. EEG data was detrended and bandpass filtered in a forward and backward direction using a 3rd-order Butterworth filter between 10-13.5 Hz for late-fast spindles, and between 14.5-18 Hz for early-fast spindles. Note that early-fast and late-fast spindles share frequency ranges near 14 Hz^20^, therefore the range of 13.5-14.5 Hz was excluded from spindle identification to reduce cross-identification of spindle event types. The upper spindle envelopes were calculated and an amplitude threshold of 75% percentile of the root mean squared value with a length window of 0.5 sec to 3.0 sec was used to define spindle events. An absolute threshold of 40 microvolts in range was used to eliminate artifacts and only spindle envelopes within 8 standard deviations from the mean amplitude values were used in further analysis. Individual slow wave events were sorted by their co-localization with each spindle type in the time domain. Spindles occurring within 0.2 sec to 1.5 sec from the trough of each slow wave were classified as a coupled event and sorted as late-fast or early-fast coupled slow wave events. Note that the time window adjacent to the slow wave troughs was buffered by 0.2 sec after the trough to avoid misidentification of slow spindles, which share frequency components with late-fast spindles and occur immediately prior to the slow wave trough.^20^ These parameters we selected to reduce mis-matching of spindles that overlap in timing and frequency components, although given the restrictive quality of these parameters, a portion of slow wave events will necessarily remain unclassified.

### Statistical Analysis

All statistical modeling was performed with SAS v9.4 (SAS Institute Inc., Cary, NC, USA).

#### Cross-channel comparison of SWA event count composition

Given that slow wave source localization experiments report both left frontal and central sources of slow waves^13^, we analyzed whether the occurrence of slow wave/spindle subtypes differs between left frontal versus central recording locations. Specifically, to compare the occurrence of late-fast and early-fast slow wave/spindle events, counts of each event type across channels (CZ and FP1) were analyzed with negative binomial generalized estimating equations (GEE).

#### Cross-channel comparison of SWA ROI spectral composition

The normalized spectral power within frontal and central channels were further compared within the ROIs corresponding to early-fast and late-fast slow wave/spindle events. Here, repeated measures regression models were used to analyze ROI data across channels (CZ and FP1).

#### Comparison of sleep stage-specific SWA composition

The SWA composition of early-fast spindle/slow wave and late-fast spindle/slow wave event counts and percentage of late-fast spindle/slow wave were measured in stages N2 and N3 sleep for each participant to quantify the effect of shifting SWA composition between cycles of NREM sleep. We performed two-sample, two-sided T-tests of each channel configuration used to detect slow wave and spindle events to analyze differences in event counts and percentage of late-fast spindle slow wave events between sleep stages (given differences in the number of slow wave/spindle events detected, calculations were weighted by the number of detected events for each participant to improve statistical reliability).

#### Comparison of SWA event count composition between age groups

Prior work has demonstrated differences in early fast and late fast slow wave/spindle EEG power^20^, and here we sought to measure changes in the composition of SWA slow wave/spindle subtypes that may correspond with these age-related EEG power changes. To measure the composition of SWA, the ratio of early fast and late fast slow wave/spindle coupling types were compared between 18–35-year-old patients and 60-70 patients with a two-sample, two-sided T-test (similar to prior analyses, calculations were weighted by the number of detected events for each participant to improve statistical reliability).

#### Comparison of SWA event percentage composition between age groups

As an additional means to measure differences in the composition of SWA among younger and older individuals, we examined regions of the overnight recordings that were heavily composed of late-fast slow wave/spindle subtypes by counting the number of time bins where the SWA composite was 80% or more of the late-fast slow wave/spindle subtype. The count of bins greater than 80% were compared between 18–35-year-old patients and 60-70-year-old patients with a negative binomial count model.

## Results

### Coupling of slow waves to spindles demonstrates an interspersed composite of distinct slow wave/spindle events (DREAMS dataset)

To aid in conceptualization of the composite structure of SWA, we identified each subtype of slow wave/spindle pair within the filtered EEG timeseries data of an individual subject (DREAMS Subject 3, see Table 1). Segments of EEG data, each with decrementing time durations, are illustrated alongside superimposed markers for each identified slow wave/spindle event to visually demonstrate the intermixed amalgam of SWA (Figure 1).

**Figure 1.**
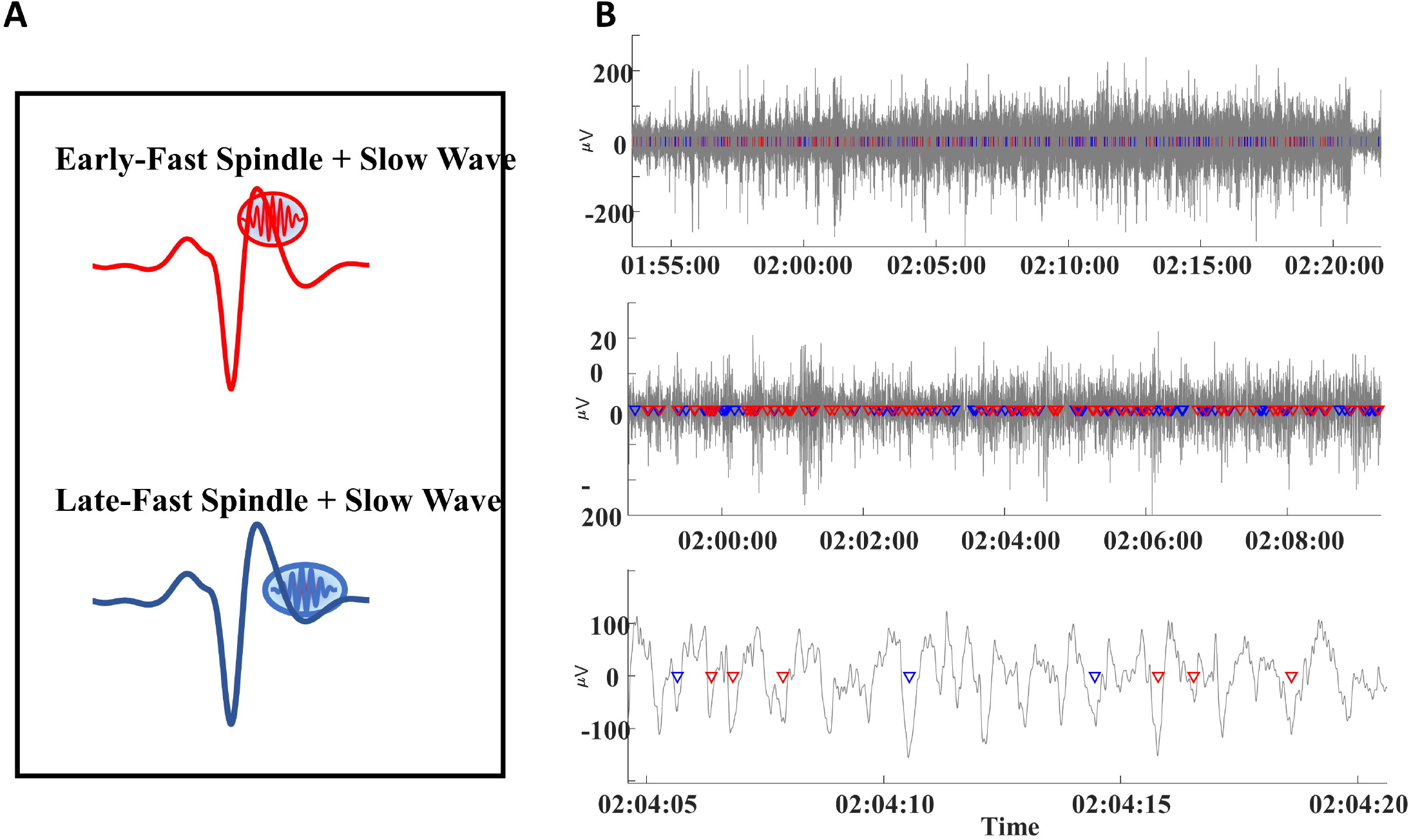
(A) Early-fast and late-fast spindle/slow wave couples are illustrated and color-coded as red and blue, respectively. (B) EEG time-series data are graphed at three levels of time duration to demonstrate the intermixing of early-fast spindle/slow waves (red markers) and late-fast spindle/slow waves (blue markers).

### Subtypes of slow wave/spindle couples are differentially represented in frontal versus central electrode locations (DREAMS dataset)

We hypothesized that frontal slow waves may demonstrate distinct coupling patterns, compared to central slow waves, and we examined slow wave coupling from channels FP1 (left frontal) and CZ (central) to test our hypothesis. Total numbers of slow waves identified were not significantly different between frontal versus central recording sites (p > 0.727), although late-fast spindle counts were ~14.8% higher in the frontal channel (95% CI: (10.9%, 18.9%), p<0.0001) compared to the central channel, and early-fast spindle counts were ~11.7% higher in the central channel compared to the frontal channel (95% CI: (7.7%, 15.8%), p=0.0004; Table 2). Event counts for slow waves coupled to late-fast versus early-fast spindles similarly demonstrated a frontal versus central preference, with late-fast/slow wave coupled events ~18.3% higher in the frontal channel (95% CI: (11.9%, 25.1%), p=0.0003), and early-fast/slow wave coupled events ~20.8% higher in the central channel (95% CI: (13.7%, 28.4%), p=0.0003). Together, these data indicate that although the number of slow waves identified in channels FP1 and CZ is similar, the mix of different slow wave/spindle couples differs between frontal versus central recording locations (Table 2).

**Table 2.** Slow wave and spindle event counts from channels FP1 and CZ among the DREAMS subjects.

| **EEG Channel** | **Mean Total**<br>**Slow Waves**<br>**(*SD*)** | **Mean Early-**<br>**Fast Spindles**<br>**(*SD*)** | **Mean Late-**<br>**Fast Spindles**<br>**(*SD*)** | **Mean Coupled**<br>**Early-Fast**<br>**+Slow Waves**<br>**(*SD*)** | **Mean Coupled**<br>**Late-Fast**<br>**+Slow Waves**<br>**(*SD*)** |
| --- | --- | --- | --- | --- | --- |
| **FP1** | 7289.30<br>(1286.37) | 2608.25<br>(649.68) | 3192.05<br>(629.39) | 1400.60<br>(462.25) | 1977.95<br>(448.41) |
| **CZ** | 7267.10<br>(1323.57) | 2912.35<br>(717.88) | 2780.00<br>(610.75) | 1692.10<br>(516.90) | 1671.90<br>(436.68) |

In addition to these differences in event counts, we also observed differences in the averaged EEG power of coupled spindles between frontal and central sites. Congruent with event count data, the normalized, averaged EEG power is ~6.5% higher for coupled late-fast spindles in the frontal region (95% CI: (3.4%, 9.5%), p=0.0002), while the normalized, averaged early-fast coupled spindle power is ~6.2% higher in the central region (95% CI: (1.8%, 10.6%), p=0.0079; Figure 2; Table 2).

**Figure 2.**
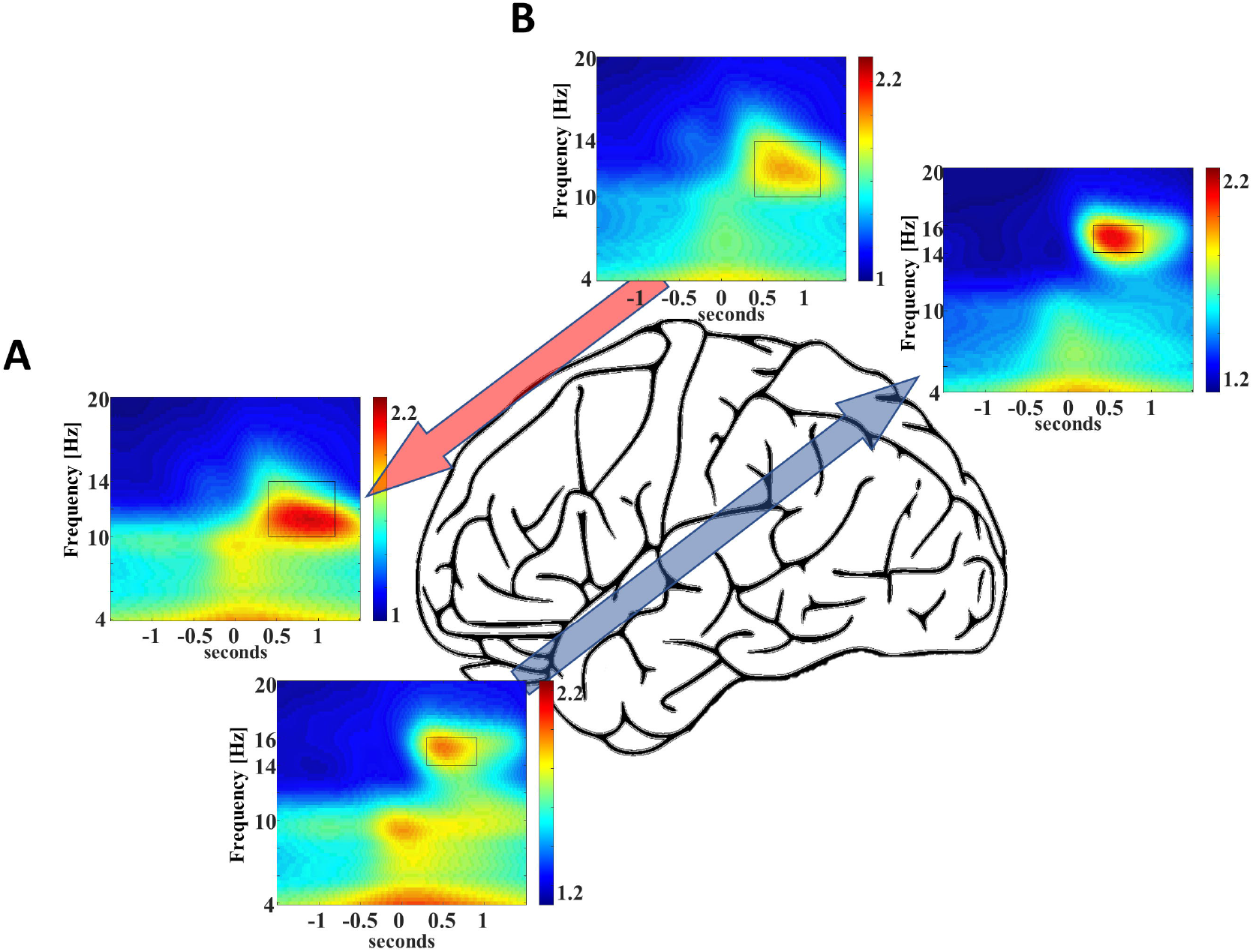
Frontal recording site FP1 demonstrates higher late-fast spindle EEG power (A), while central recording site Cz demonstrates higher early-fast spindle EEG power (B).

### The composite of SWA is formed by a mixture of slow wave/spindle coupled events, and their relative contribution to the SWA rises and falls across cycles of NREM sleep (DREAMS dataset)

In consideration of recent studies that demonstrate each instance of slow wave/spindle coupled event produces a cortical reinstatement of recent memory representations^16,38^, we sought to measure the instances of each slow wave/spindle event subtype across cycles of NREM sleep. Congruent with measures of coupled spindle power, there were distinctions between stages N2 and N3 in the composition of slow wave/spindle subtypes. At the level of the individual, each cycle of NREM sleep is observed to shift in the composition of SWA, and a distinction between production of late-fast versus early-fast coupled slow waves moves through the night in accordance with the with the depth of sleep (illustrated for a single participant in Figure 3).

**Figure 3.**
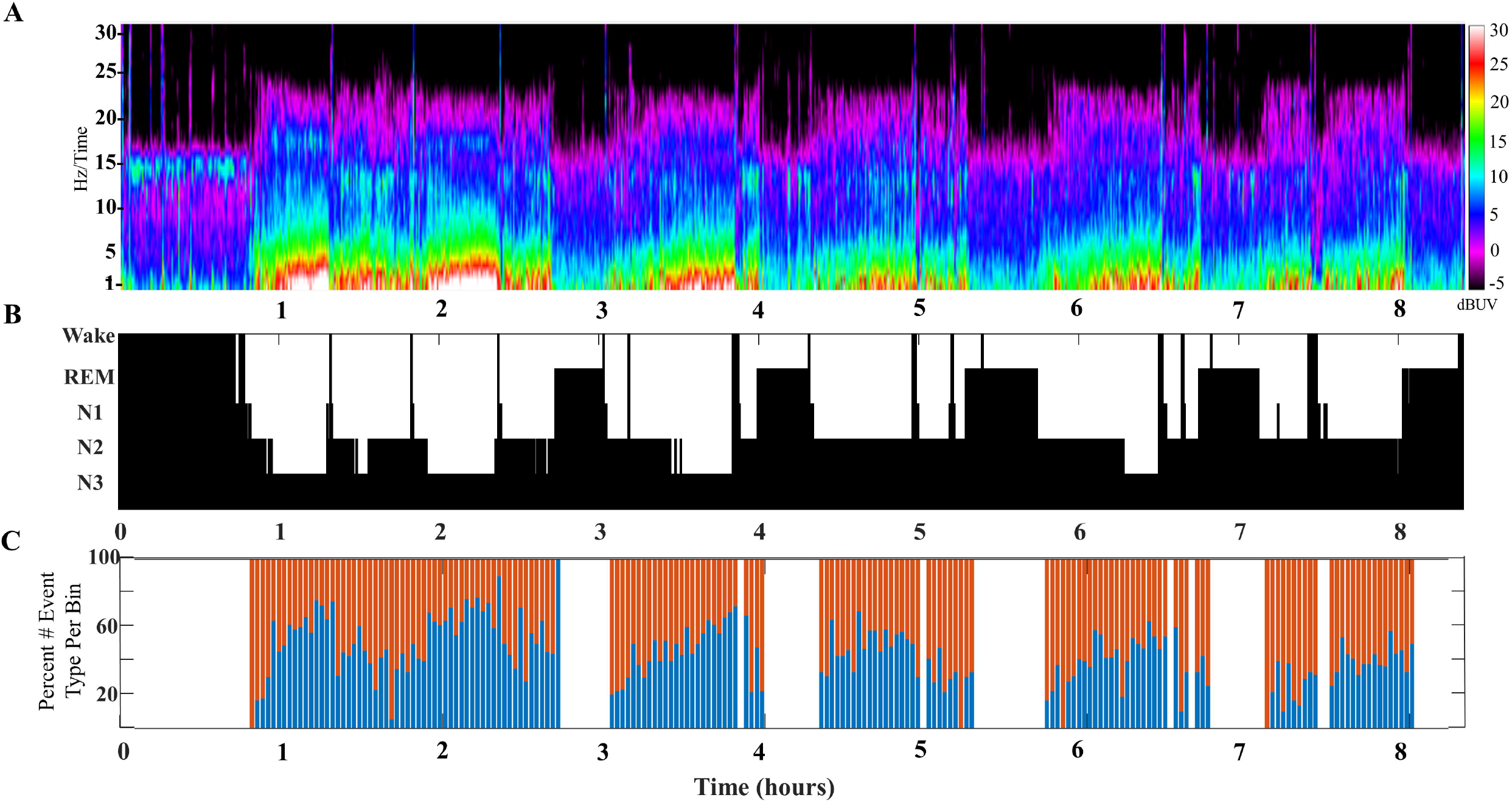
(A) An illustrative example of SWA composition for a single participant aged 24 years old (DREAMS Subject 3). A color density spectral array provides a graphic illustration of EEG power for an entire overnight sleep recording of a single subject. (B) The hypnogram marking sleep stages for this subject’s overnight sleep recording is illustrated. (C) The percentage of early-fast spindle/slow waves and late-fast spindle/slow waves in each time bin of 2 minutes is graphed to illustrate the composite mix of SWA across a single night of sleep.

This undulation of SWA composite through NREM cycles is most clearly observed when slow wave/spindle subtypes are identified with respect to their location propensity, i.e., frontal for late-fast and central for early-fast coupled slow wave/spindles. Identification of both subtypes from the same channel location yielded similar results, although the shifts between SWA composition were less pronounced (illustrated in Figure 4).

**Figure 4.**
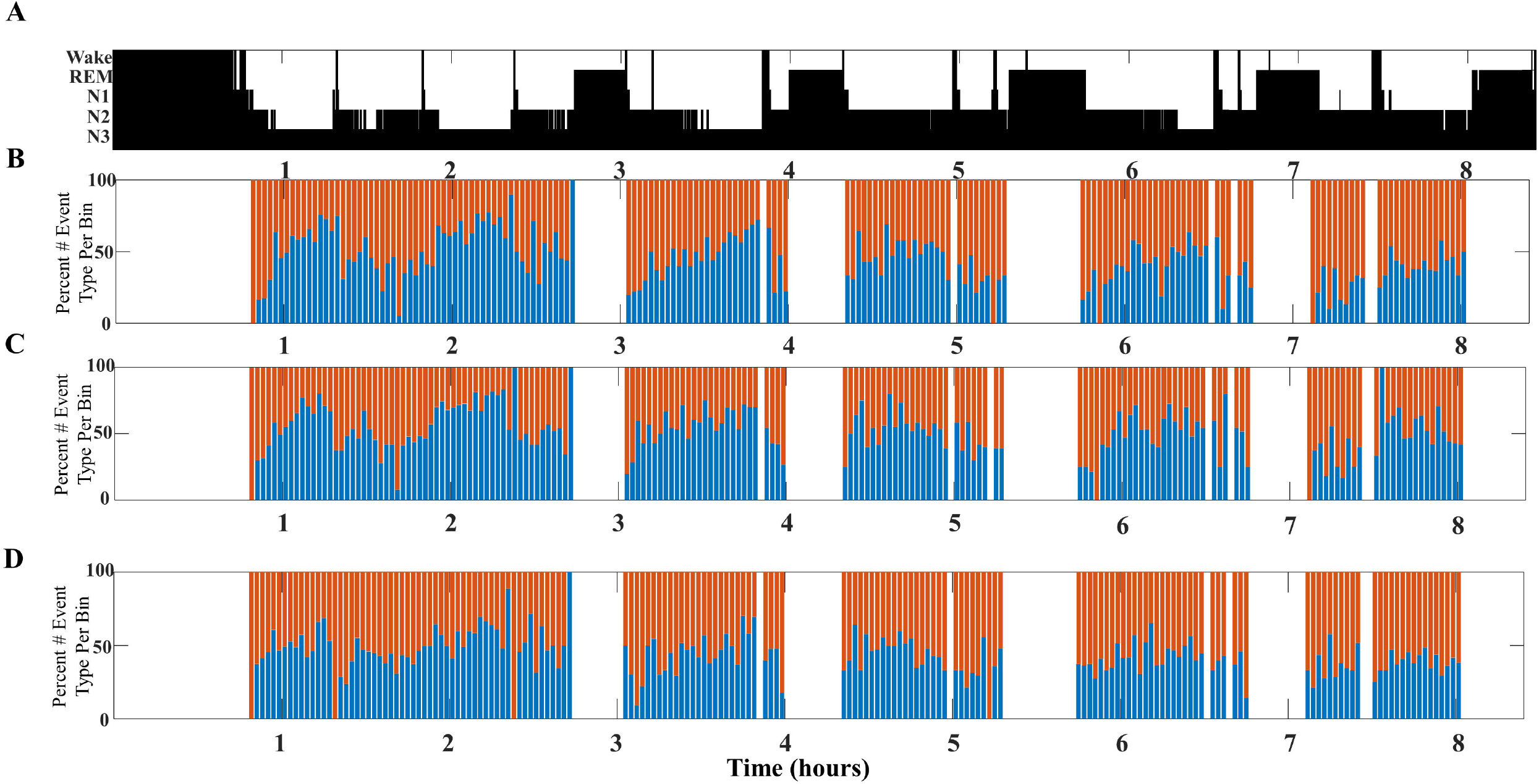
An illustrative example comparing SWA composition as measured from different channel pairs for a single participant (DREAMS Subject 3). (A) A hypnogram marking sleep stages for a single participant’s overnight sleep recording is illustrated. (B) The percentage of early-fast spindle/slow waves (red) detected in channel CZ and late-fast spindle/slow waves (blue) detected in channel FP1 are graphed in time bins of 2 minutes. (C) The percentage of early-fast spindle/slow waves (red) detected in channel FP1 and late-fast spindle/slow waves (blue) detected in channel FP1 are graphed in time bins of 2 minutes. (D) The percentage of early-fast spindle/slow waves (red) detected in channel CZ and late-fast spindle/slow waves (blue) detected in channel CZ are graphed in time bins of 2 minutes.

To further illustrate the shifts in SWA composition across individual subjects’ NREM sleep cycles, eight additional participants from the DREAMS database are presented with event histograms marking normalized percentage of total stage N2 and N3 sleep for each individual (Figure 5).

**Figure 5.**
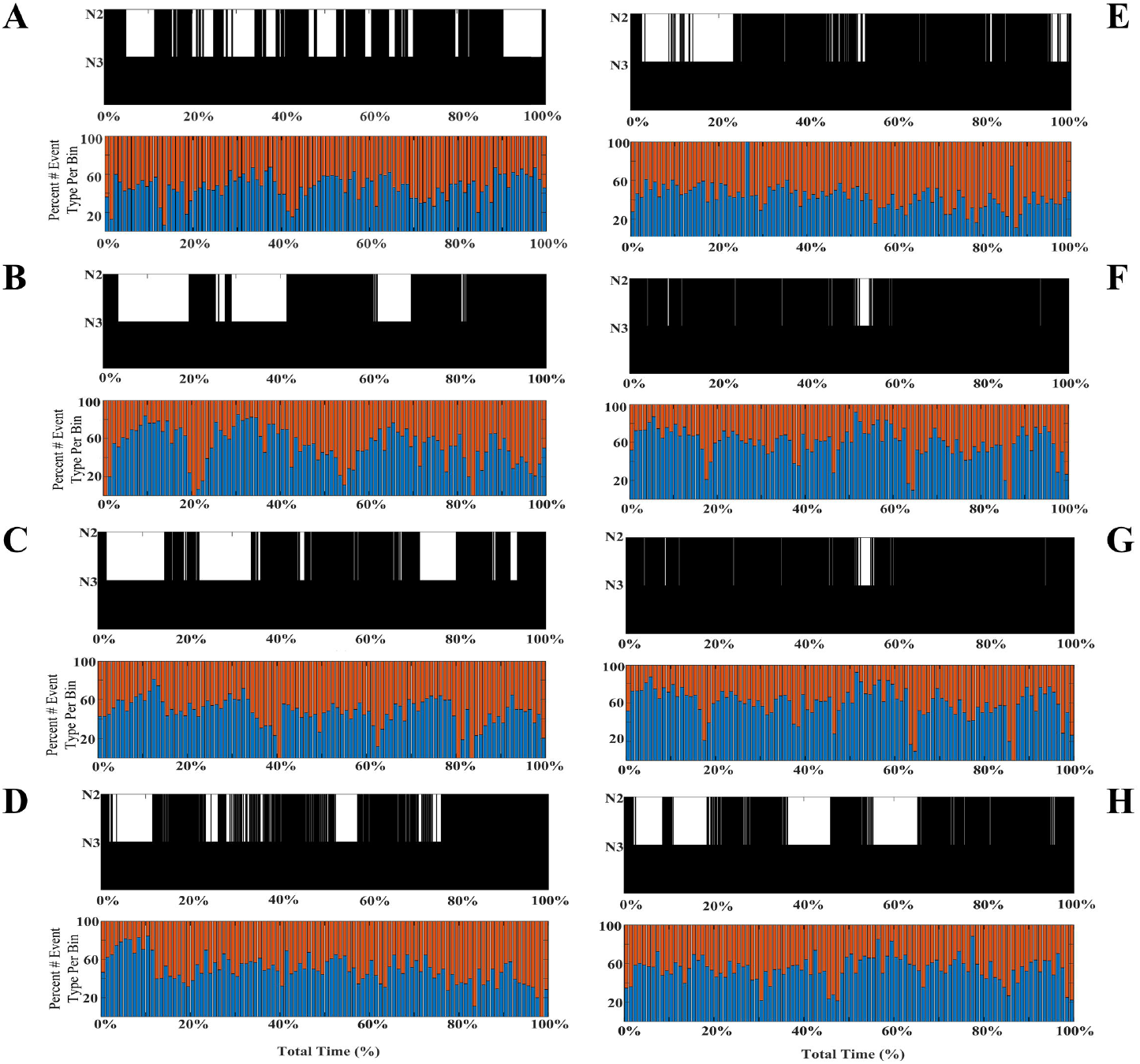
Individual differences in the shifting composition through stages N2 and N3 SWA. Event histograms marking the percentage of early-fast spindle/slow waves detected in channel CZ (red) and late-fast spindle/slow waves detected in channel FP1 (blue) are graphed in time bins of 2 minutes (A-H; DREAMS Subjects 1, 2, 4-9).

### Quantification of SWA composition between stages N2 and N3 sleep confirms stage-specific composition shifts underly undulations across cycles of NREM sleep (DREAMS dataset)

Building on the observation that SWA composition shifts across cycles of NREM sleep, we next measured the number of each slow wave/spindle subtype in each stage of sleep from all 20 participant recordings in the DREAMS database to calculate the percentage of late-fast versus early-fast coupled slow waves for N2 and N3 sleep. Our visual representations of SWA composition suggest that the undulations of SWA shift toward greater late-fast slow wave/spindle subtypes in N3 sleep, while early-fast slow wave/spindle subtypes are more frequent in stage N2, congruent with prior observations that demonstrate a distinction in the late-fast spindle and early-fast spindle EEG power between stages N2 and N3 sleep^20^. Quantifying the percentage of late-fast versus early-fast slow wave/spindle events from stages N2 and N3 sleep was congruent with this observation. Measuring late-fast slow wave/spindle events from channel FP1 and early-fast slow wave/spindle events from channel CZ, 59.93% (95% CI: (55.95%, 63.92%)) of slow wave/spindle events during N3 sleep were late-fast slow wave/spindle subtype, while only 49.25% (95% CI: (46.12%, 52.39%)) of N2 sleep were the late-fast subtype (FP1-CZ pair, estimated difference = 10.68 percent units, 95% CI: (6.78, 14.57), p<0.0001; Table 3). Measuring both late-fast slow wave/spindle events and early-fast slow wave/spindle events from channel CZ, 53.15% (95% CI: (48.36%, 57.95%)) of slow wave/spindle events during N3 sleep were late-fast slow wave/spindle subtype, while only 48.08% (95% CI: (44.70%, 51.46%)) of N2 sleep were the late-fast subtype (CZ-CZ pair, estimated difference = 5.08 percent units, 95% CI: (1.33, 8.82), p=0.0114; Table 3). Finally, measuring both late-fast slow wave/spindle events and early-fast slow wave/spindle events from channel FP1, 64.99% (95% CI: (60.88%, 69.11%)) of slow wave/spindle events during N3 sleep were late-fast slow wave/spindle subtype, while only 53.04% (95% CI: (50.04%, 56.03%)) of N2 sleep were the late-fast subtype (FP1-FP1 pair, estimated difference = 11.96 percent units, 95% CI: (7.66, 16.26), p<0.0001; Table 3). We further performed a comparative analysis of the differences in SWA composition between these channel pairs (FP1-CZ, CZ-CZ, and FP1-FP1). Additional statistical comparisons between channel pairs (FP1-CZ, CZ-CZ, and FP1-CZ) were performed for percentage of late-fast spindle/slow wave pairs in both stages N2 and N3 sleep. Each permutation of channel comparison (i.e., FP1-CZ to CZ-CZ and FP1-FP1; CZ-CZ to FP1-CZ and FP1-FP1; FP1-FP1 to FP1-CZ and CZ-CZ) were statically significant (p<0.001) for each comparison, except for late-fast spindle/slow percentage in FP1-CZ versus CZ-CZ (p=0.06; Table 3).

**Table 3:** Stage N3- and N2-specific slow wave and spindle event counts and percentage of late-fast slow wave/spindle pairs from each channel pairing among the DREAMS subjects

| **EEG Channel Pair** | **N3 Sleep Mean Coupled Early-Fast +Slow Waves (SD)** | **N3 Sleep Mean Coupled Late-Fast +Slow Waves (SD)** | **N2 Sleep Mean Coupled Early-Fast +Slow Waves (SD)** | **N2 Sleep Mean Coupled Late-Fast +Slow Waves (SD)** | **N3 Sleep Mean Percent Late-Fast +Slow Waves (SD)** | **N2 Sleep Mean Percent Late-Fast +Slow Waves (SD)** |
| --- | --- | --- | --- | --- | --- | --- |
| **FP1-CZ** | 736.95<br>(389.56) | 1058.50<br>(435.30) | 955.15<br>(285.72) | 919.45<br>(382.03) | 59.90<br>(9.57) | 48.55<br>(6.22) |
| **CZ-CZ** | 736.95<br>(389.56) | 796.05<br>(345.28) | 955.15<br>(285.72) | 875.85<br>(351.63) | 53.14<br>(11.80) | 47.53<br>(6.73) |
| **FP1-FP1** | 583.65<br>(352.12) | 1058.50<br>(435.30) | 816.95<br>(259.25) | 919.45<br>(382.03) | 65.56<br>(9.27) | 52.49<br>(5.54) |

| **EEG Channel Pair** | **N3 Sleep Mean Weighted Percent Late-Fast +Slow Waves (SD)** | **N2 Sleep Mean Weighted Percent Late-Fast +Slow Waves (SD)** |
| --- | --- | --- |
| **FP1-CZ** | 58.95<br>(8.76) | 49.05<br>(6.88) |
| **CZ-CZ** | 51.93<br>(10.66) | 47.83<br>(7.42) |
| **FP1-FP1** | 64.46<br>(8.90) | 52.95<br>(6.58) |

### The composition of SWA among older adults favors a higher proportion of late-fast coupled slow wave/spindle events, and higher percentages of late-fast spindles during peak times (CFS dataset)

To examine how the composition of SWA differs with age, we compared the slow wave/spindle subtype structure of NREM cycles between young adults and older adults from the CSF dataset. Overall, the mean proportion of late-fast slow wave/spindle coupled events across the entire night is 57.1% (95% CI: (54.9%, 59.3%)) in older adults, compared to 53.7% (95% CI: (52.0%, 55.4%)) in younger adults (estimated difference = 3.43 percent units, 95% CI: (−0.7%, −6.2%), p=0.0146; Table 4). Further, the peaks of high late-fast spindle/slow wave SWA composition frequently demonstrate a substantially higher amount of late-fast spindle coupling, when compared to the highest peaks of late-fast SWA among young adults (illustrated in Figure 6). To quantify this observation, a threshold was set to 80% normalized late-fast slow wave/spindle events per 1% of the total normalized NREM time for each subject, and the number of bins exceeding this threshold was quantified among the young adults and older adults (Figure 6). Older adults averaged 10.9 (95% CI: (8.5, 14.0)) bins greater than or equal to 80% late-fast slow wave/spindle events, while young adults averaged 2.9 (95% CI: (2.0, 4.2)) of these high late-fast bins (older versus younger ratio = 3.76, 95% CI: (2.41, 5.87), p<0.0001; Table 4).

**Figure 6.**
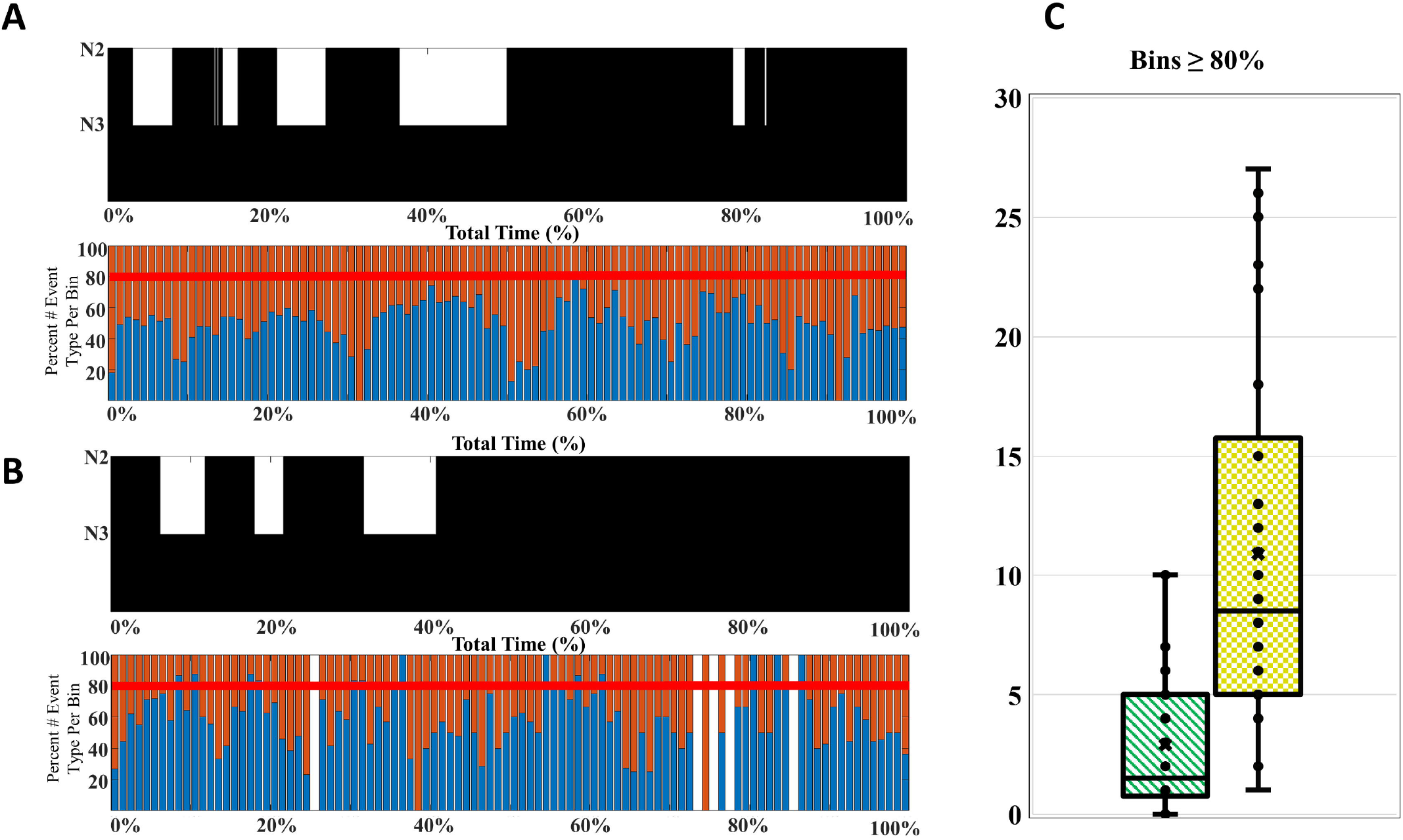
A comparison of the SWA composition among younger and older adults. (A) A younger participant (CFS 800092), age 21 y.o., and (B) an older participant (CFS 800099), age 61 y.o., are presented as an illustrative example of SWA thresholding. The percentage of late-fast spindle/slow wave events (blue) and early-fast spindle/slow wave events (red) are assessed for bins with 80% or greater late-fast spindle/slow wave composition. (C) The number of bins at or exceeding 80% late-fast spindle/slow wave composition for each participant is quantified, and the younger adults (green-stripes; n=30) are compared to older adults (yellow-checkered; n=30) at the group level.

**Table 4.** Slow wave and spindle event counts, and mean number of bins exceeding a threshold of 80% composition of the late-fast subtype, for younger and older adults from the Cleveland Family Study.

| **Age Group** | **Mean Total Slow Waves (SD)** | **Mean Early-Fast Spindles (SD)** | **Mean Late-Fast Spindles (SD)** | **Mean Coupled Early-Fast+Slow Waves (SD)** | **Mean Coupled Late-Fast+Slow Waves (SD)** | **Mean Number of Bins $\geq 80\%$ (SD)** |
| --- | --- | --- | --- | --- | --- | --- |
| **Younger Adults** | 5128.67<br>(1556.93) | 3020.23<br>(736.80) | 3355.67<br>(792.60) | 1176.93<br>(422.75) | 1363.43<br>(500.76) | 2.9<br>(3.03) |
| **Older Adults** | 4111.10<br>(1145.40) | 2198.43<br>(455.41) | 2404.00<br>(568.77) | 757.97<br>(244.63) | 1008.93<br>(360.52) | 10.9<br>(7.79) |

## Discussion

Our analyses demonstrate that SWA can be decomposed into distinct subtypes based on coupling of slow wave and spindle pairs. Results from the study suggest that each subtype of slow wave/spindle event demonstrates unique neurophysiological properties that predict their rate of occurrence, including electrode location bias and relative depth of sleep. Collectively our findings build upon and may provide increased clarity on prior work that demonstrates slow waves have anterior and central sources, and that SWA differs in cortical propagation and homeostatic properties along the anterior-to-posterior axis.

Our whole-night illustration of SWA composition further demonstrate a striking rise and fall of slow wave/spindle events across cycles of NREM sleep, suggesting the possibility of competing or complementary processes that occur in a choreographed rise and fall of SWA composition states. In addition, our results demonstrate that the event composition of SWA is quantitatively distinct between stages N2 and N3 sleep, with deeper sleep favoring a higher percentage of late-fast slow wave/spindle events. This is consistent with our prior observations that demonstrate the normalized EEG power differs in the late fast versus early fast ROI regions within slow wave/spindle events^20^, and suggests that the differences in EEG power may be, at least in part, attributable to differences in the number of each slow wave/spindle subtype that creates each sleep stage’s SWA composite.

Also congruent with our previous analyses of slow wave/spindle EEG power,^20^ we observed that aging is associated with a relative loss of early-fast slow wave/spindle events, as the late-fast coupled slow waves form a greater contribution to the mix of SWA and increasingly dominate the SWA event composition. Here we further demonstrate that extremes of SWA event composition are common among aging adults, and late-fast coupled slow wave/spindle pairs dominate these segments of skewed SWA composition.

Understanding that SWA is not uniform, but instead shifts in composition across cycles of NREM, provides key insights to inform interpretations of prior studies and better design future experiments. One such scenario may be studies of slow wave coupling and memory performance, which have reported seemingly conflicting relationships^4,23^. Key differences in experimental design, particularly regarding the captured and analyzed depth of slow wave sleep, may be reflected in the composition of SWA among study participants and may account for discrepancies in outcome measures. Further, experimental manipulation of SWA may preferentially impact one type of slow wave/spindle event, creating an imbalance in the composite of SWA across portions of NREM sleep. Accounting for these changes in slow wave physiology may be critical to understanding the mechanisms underlying SWA manipulation.

While the function of early-fast versus late-fast coupling with slow waves is yet to be determined, additional mechanistic insights may come from intracranial recording techniques. Human experiments among patients undergoing epilepsy surgery have described distinctions between anterior and posterior hippocampal slow wave/spindle coupling^18,19^. Given that the anterior and posterior hippocampi are proposed to utilize distinct neuroanatomical connections for processing different forms of memory content^40,41^, slow wave/spindle coupled memory processing events may also reflect different forms of memory processing in frontal versus parietal cortical regions. Indeed, data from experimental manipulation of SWA in an animal model demonstrate two distinct forms of slow wave/spindle coupling, and their functions may be opposing to one another in synaptic regulation and memory processing functions^21^.

Our analyses delineate key neurophysiological properties of late-fast and early-fast spindle coupling, although important limitations remain in characterizing the composition of SWA and the function of each distinct type of slow wave and spindle event. Signal processing experiments have utilized features of slow waves including the waveform characteristics and recording locations to describe multiple subtypes of slow waves as anterior versus central^13^, global versus local^12^, and fast switching versus slow switching (trough-to-peak transition frequency)^42^. In addition, the coupling of slow waves with spindle and hippocampal ripples is reported to differ between anterior versus posterior hippocampal locations, further suggesting that a division of coupled slow waves may follow an similar anterior versus posterior pattern.^18,19^ Future experiments will be required to determine whether spindle coupling indeed separates slow wave events into these previously described slow wave subtypes. In addition, the use of coupling to identify slow waves has inherent limitations in the ability to label slow wave events, including an overlapping frequency component that precludes characterization of many spindles in this shared frequency range at approximately 14 Hz^20^. Further, many slow waves do not clearly demonstrate a coupled spindle on surface EEG, and there remain numerous undifferentiated slow waves that may represent additional subtypes of slow wave events. The incorporation of additional slow wave and spindle features may provide the next steps in better decomposing all SWA in future studies.

Aging has been linked to changes in slow wave/spindle coupling^26,35^, and indeed, the continuum of the adult human lifespan demonstrates a shift in the composition of coupled spindle EEG power towards the late-fast subtype^20^. The results of this study support a conceptual model of a drift in the types of slow wave/spindle events that older adults produce, and a striking increase in the number of time periods that cycle disproportionately into late-fast spindle domination of SWA. Given the interest in spindle coupling as a biomarker of cognitive aging and risk of neurodegenerative disease^7,8,24,28^, the potential impact of this late-fast spindle coupling bias among aging adults may provide mechanistic insights and facilitate the development of a predictive functional biomarker. Future studies may focus on the neuroanatomical basis of late-fast and early fast spindle to better understand the potential mechanisms that may produce a shift in composition of sleep.

## Acknowledgements

This work was supported in part by funding from the National Institutes of Health (R01AG058772). The authors are also grateful for the open access to data provided by the DREAMS database (10.5281/zenodo.2650142) and the Cleveland Family Study (CFS) (supported by grants from the National Institutes of Health (HL46380, M01 RR00080-39, T32-HL07567, RO1-46380)), and the National Sleep Research Resource (supported by the National Heart, Lung, and Blood Institute (R24 HL114473, RFP 75N92019R002)).

## Author Contributions

B.V.M., E.K., V.O.K., S.H.S., A.R.R, R.L.P. and B.M.B. designed the study; B.V.M., S.H.S, L.M.M., and R.L.P. analyzed the data; B.V.M. and B.M.B. wrote the manuscript; E.K., V.O.K., and A.R.R. gave conceptual advice. All authors participated in editing and revision of the manuscript.

## Financial Disclosure

None

## Non-financial disclosure

None

*The authors declare that the research was conducted in the absence of any commercial or financial relationships that could be construed as a potential conflict of interest.*

The data underlying this article are available in the DREAMS database at https://doi.org/10.5281/zenodo.2650142/, and the National Sleep Research Resource, at https://sleepdata.org/.

## Notes

### Competing Interest Statement

The authors have declared no competing interest.

